# How the Structure of Signaling Regulation Evolves: Insights from an Evolutionary Model

**DOI:** 10.1101/2024.10.23.619883

**Authors:** Danial Asgari, Ann T. Tate

## Abstract

To remain responsive to environmental changes, signaling pathways attenuate their activity with negative feedback loops (NFLs), where proteins produced upon stimulation downregulate the response. NFLs function both upstream of signaling to reduce input and downstream to reduce output. Unlike upstream NFLs, downstream NFLs directly regulate gene expression without the involvement of intermediate proteins. Thus, we hypothesized that downstream NFLs evolve under more stringent selection than upstream NFLs. Indeed, genes encoding downstream NFLs exhibit a slower evolutionary rate than upstream genes. Such differences in selective pressures could result in the robust evolution of downstream NFLs while making the evolution of upstream NFLs more sensitive to changes in signaling proteins and stimuli. Here, we test these assumptions within the context of immune signaling. Our minimal model of immune signaling predicts robust evolution of downstream NFLs to changes in model parameters. This is consistent with their critical role in regulating signaling and the conservative rate of evolution. Furthermore, we show that the number of signaling steps needed to activate a downstream NFL is influenced by the cost of signaling. Our model predicts that upstream NFLs are more likely to evolve under a shorter half-life of signaling proteins, absence of host-pathogen co-evolution, and a high infection rate. Although it has been proposed that NFLs evolve to reduce the cost of signaling, we show that a high cost does not necessarily predict the evolution of upstream NFLs. The insights from our model have broad implications for understanding the evolution of regulatory mechanisms across signaling pathways.

## Introduction

Signaling pathways that respond to external stimuli are tightly regulated to maintain homeostasis. The external signal is typically received by upstream proteins that are located near the cell surface and then passed on to downstream proteins to activate target genes. Gene expression during signaling can impose energetic costs and excessive expression can disrupt cellular functions (Dekel & Alon, 2005). To prevent this, signaling is regulated by negative feedback loops (NFLs). Upon stimulation, NFLs are activated to attenuate signaling by downregulating both upstream and downstream proteins, thereby regulating the response at multiple levels within signaling pathways (Nair et al., 2019). NFLs that regulate signaling by targeting receptors and other upstream proteins dampen the response by decreasing input. On the other hand, NFLs that act on transcription factors and other downstream proteins, reduce gene expression while maintaining signaling from upstream components. Therefore, unlike upstream NFLs, downstream NFLs directly regulate gene expression in response to stimuli. This can impose different selective pressures on downstream and upstream NFLs, leading to the evolution of these NFLs under different conditions.

Multi-level regulation of signaling by upstream and downstream NFLs appears to be evolutionarily conserved across biological networks. Wnt signaling, for example, which is present in all multicellular animals, is regulated by an upstream NFL involving DKK, which reduces input into the pathway via receptor endocytosis (J. Liu et al., 2022). On the other hand, Axin2 acts downstream of Wnt signaling to negatively regulate gene expression by inhibiting β-catenin (Lustig et al., 2002). Other conserved biological pathways are also regulated at different levels of signaling. The mTOR signaling that regulates fundamental cellular functions such as metabolism, growth, and survival is modulated by mTORC1 and S6K, which are NFLs that negatively regulate the upstream insulin receptor substrate (Rozengurt et al., 2014). Hippo signaling, which has various biological functions and is linked to many cancerous and non-cancerous diseases, is regulated by an NFL involving LATS1/2 that targets the downstream protein complex YAP/TAZ to reduce gene expression (Dai et al., 2015). These examples show the prevalence of multi-level regulation by NFLs within diverse signaling pathways and species.

Previous models studied NFLs to understand how the positioning of NFLs and their interaction with positive feedback loops shape the response to stimuli. For example, Pfeuty and Kaneko (2009) have shown that NFLs are necessary for switching between stable states induced by positive feedback loops. Another study has shown that the positioning of multiple NFLs within signaling pathways can either induce or abolish oscillations (Nguyen, 2012). These studies shed light on the role of feedback loops in the dynamics of singling. However, without understanding the evolutionary forces that act upon upstream and downstream NFLs, many questions regarding the diversity and function of NFLs remain unanswered. For example, why do some upstream NFLs directly interact with receptors whereas others interact with the stimulator that activates the receptor? Why do some downstream NFLs show remarkable consistency in the mechanism of actions across diverse signaling pathways? Why do different signaling pathways use different numbers of upstream and downstream NFLs?

Because immune signaling pathways must contend with an evolving stimulator (i.e., pathogen) and at the same time minimize immunopathology (i.e. cost of signaling), they are uniquely suited for the study of multi-level regulation of signaling networks. Immune receptors can recognize pathogens directly (Kato et al., 2006) or indirectly by detecting conserved pathogen associated molecular patterns released by the pathogen (Kleino & Silverman, 2014) or by interacting with cytokines (Jang et al., 2006). These differences influence receptor-pathogen co-evolution which might in turn shape the NFL-receptor evolution. Upstream NFLs within immune signaling pathways show an interesting diversity in the mechanism of action. For example, within the Imd pathway of invertebrates, Pirk directly downregulates receptors, whereas A20 within NF-κB pathway of vertebrates indirectly downregulates the receptor by interacting with RIP-1 (Kleino et al., 2008; Pujari et al., 2013). Also, Imd signaling uses upstream NFLs such as PGRP-LB/SC to scavenge peptidoglycans before they activate receptors, thus reducing the input into the pathway without direct interaction with receptors (Costechareyre et al., 2016; Orlans et al., 2021). The end product of immune signaling pathways (i.e., effector proteins) not only kills pathogens but also harms host tissues at high doses (Bonneaud et al., 2003; Hanssen et al., 2004). The expression of immune genes is regulated by downstream NFLs, such as the Repressosome complex (STAT92E and AP-1) in invertebrates, IκBα in vertebrates, and JAM transcription factors in plants, which target downstream transcription factors (Kearns et al., 2006; Kim et al., 2007; Sasaki-Sekimoto et al., 2013). These NFLs show consistency in the mechanism of action by competing with transcription factors for binding to the promoter of target genes and shut down gene expression. These intriguing aspects of immune signaling have prompted us to explore conditions favoring the evolution of NFLs functioning at different signaling stages within the context of immune responses.

Here we hypothesized that because downstream NFLs are directly responsible for the regulation of gene expression upon receiving the signal, they evolve under a more stringent selection than upstream NFLs. Indeed, genes encoding downstream NFLs show a slower and more consistent rate of evolution across various signaling pathways compared to upstream genes (Fig.1). Here, we Identify the consequences of these evolutionary differences by using a minimal evolutionary model of immune signaling, which combines shared features across various immune signaling networks (Fig.2). Based on our hypothesis we predict that the evolution of upstream NFLs should be more sensitive to model parameters such as the half-life of signaling components, infection rate, and host-pathogen co-evolution. By dissecting the effect of these factors on the evolution of upstream and downstream NFLs, our study sheds light on the evolution of the ubiquitous phenomenon of multi-level regulation across various signaling networks.

**Fig 1.**
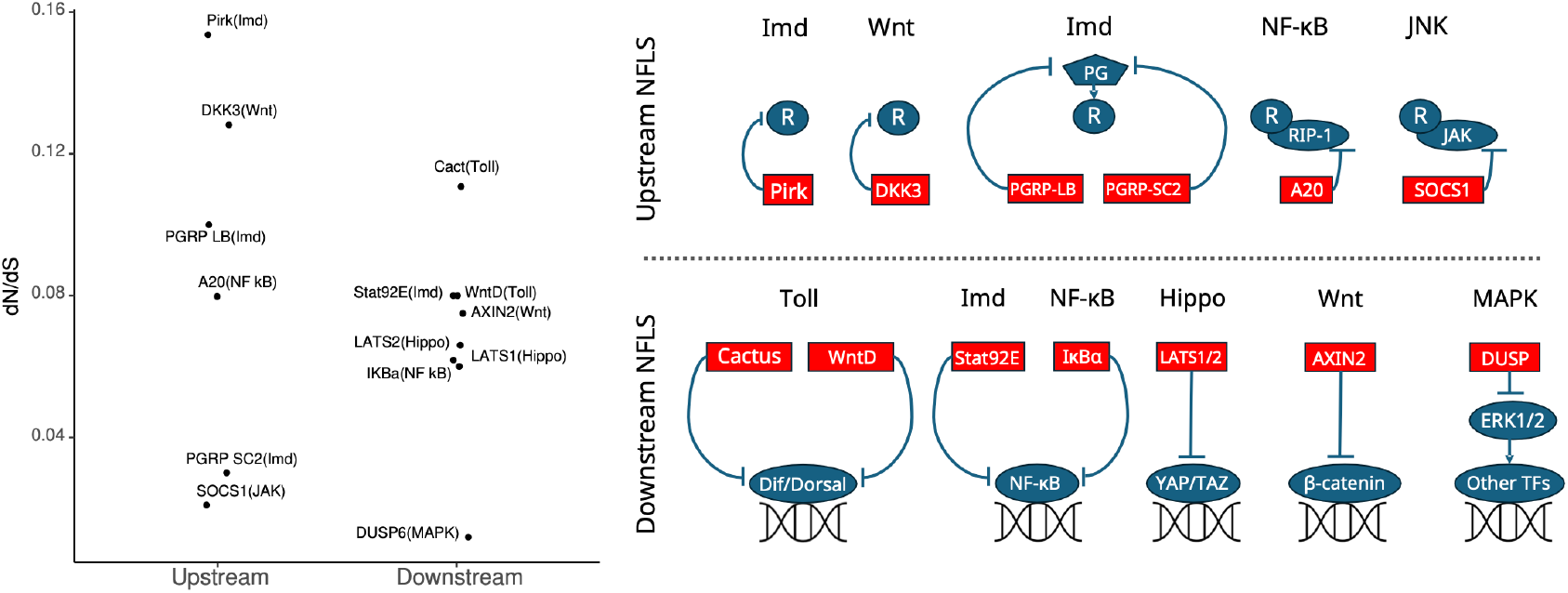
The distribution of 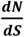 values for genes encoding proteins involved in upstream and downstream NFLs across various signaling networks. The rate of change for genes encoding upstream and downstream NFLs is shown on the left. The biological pathway is indicated in the parenthesis. The function of Upstream and downstream NFLs within various signaling pathways is summarized on the right. The upstream NFLs interact with a receptor (*R*), while downstream NFLs interact with transcription factors that activate target genes. These interactions are either direct or indirect (e.g., with peptidoglycan shown as PG). Genes involved in Imd and Toll are from *D. melanogaster* and the rest are human genes. The data for human genes are from Liu et al. (2014) except for IKB*α* which is from Song et al. (2012). The data for *D. melanogaster* is from Sackton et al. (2007) except for Pirk which is from Han et al. (2013). The downstream genes generally have low 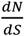 values (all except for Cactus ≤ 0.08) and the variance amongst downstream genes (*sd* = 0.028) is lower than upstream ones (*sd* = 0.053).

**Fig 2.**
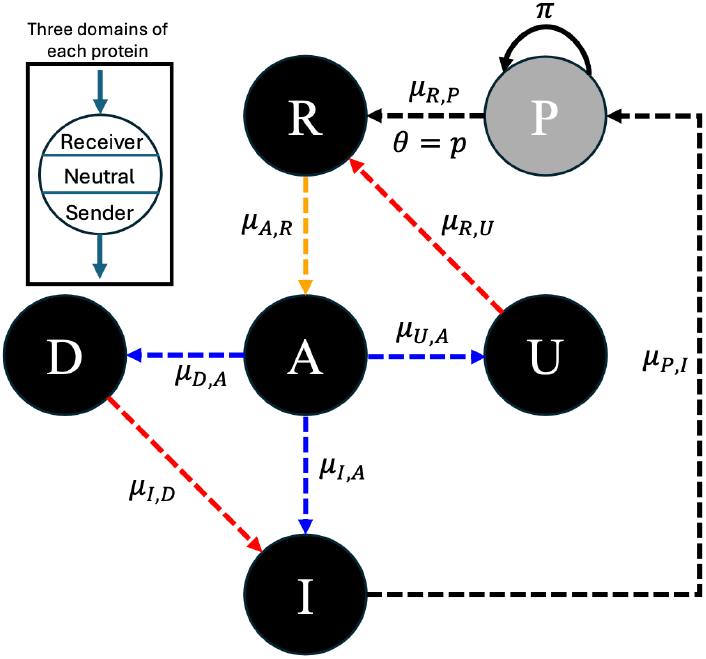
The model of the immune signaling network, capturing the core elements shared across immune signaling pathways. A host has five proteins: *R* (receptor), *A* (activator), *I* (immunity), *U* (upstream regulator), and *D* (downstream regulator). The host is exposed to a pathogen (*P*) with a probability *θ*, and the pathogen replicates at a rate *π*. The dashed arrows show potential interactions that can be formed during the evolution of the network. The arrows point from protein *j* to *i* and *μ*_*i,j*_ is the strength of the interaction. The three domains of each protein (receiver, neutral, and sender) are shown in the top right. The neutral domain does not participate in protein-protein interactions. For a detailed description of the model see Materials and Methods.

## Results

### The evolution of the downstream NFL is robust to the choice of parameters and the cost of response can influence the immune signaling architecture

To understand whether upstream and downstream NFLs evolve under distinct selective pressures (Fig.1), we used an evolutionary model of immune signaling to identify conditions under which both NFLs evolve. First, we adjusted model parameters to increase the relative cost of infection to the cost of the immune response (*α* = 2 and *β*= 1 in *Eq. 4* of Materials and Methods). This condition ensures two outcomes: the evolution of a functional immune signaling pathway that reduces pathogen burden, and that the response incurs a cost. Second, we set the degradation rates for the host proteins to 0 (*ϕ* = 0 in *Eq. 3*). This ensures that host proteins do not degrade on their own on a timescale relevant to signaling, demanding the action of NFLs to reduce the response. As we predicted, the immune signaling networks evolve to respond to the pathogen (*R* activates *A* and *A* activates *I* in Fig.3a and b); however, only the downstream NFL (*D*) evolved under these conditions (*A* activates *D* and *D* inhibits *I* in Fig.3c), while the upstream NFL (*U*) did not evolve (Fig.3d). Due to host-pathogen co-evolution, the interaction coefficients between the receptor (*R*), immunity (*I*), and pathogen (*P*) fluctuate continuously (Fig.S1).

Next, we tested the robustness of the evolution of a network with a downstream NFL by changing the degradation rate of host proteins (*ϕ*), the rate of infection (*θ*), the pathogen initial condition (*P*_0_), and its replication rate (*π*). We observed that upon changing these parameters, the downstream NFL still evolved, while the upstream NFLs did not (Fig.S2). Thus, the direct interaction between the downstream regulator (*D*) and the output (*I*) leads to a more robust evolution of the downstream NFL, while the evolution of stable interactions between the receptor and the upstream NFL was not favored under any of these conditions.

The absence of the upstream NFL amongst evolved signaling pathways under previous conditions was unexpected. We hypothesized that a low cost of immunity in previous simulations hinders the evolution of upstream NFLs. To test this, we increased the cost of the immune response (*α* = 2 and *β*= 2 in *Eq. 4* and *ϕ* = 0 in *Eq. 3*) but kept other parameter values identical to those in Fig.3. Contrary to expectations the upstream NFL still did not evolve. Furthermore, in two out of 10 representative simulations, the receptor (*R*) did not activate the activator (*A*) protein (grey trajectories in Fig.4a); thus, a functional signaling network did not evolve. Amongst the other eight simulations, two were similar to the previously evolved networks (Fig.3) where the downstream regulator (*D*) acts as an NFL to reduce the immune response (*I*), which is activated by the activator protein (*A*) (Fig.4b and c1). In six simulations, the activator (*A*) evolved as an NFL to inhibit immunity (*I*), while the downstream regulator (*D*) evolved to activate the immune response (*I*) (Fig.4b and c2). When we set *ϕ* > 0, in six simulations *A* evolved as NFL, while *D* evolved as an NFL in the remaining four (Fig.S3).

We next repeated the simulation after increasing the population size (*n*) from 2,000 to 5,000. Upon increasing the population size, the high cost of immunity (*β*= 2) resulted in the evolution of *A* as an NFL in all simulations (Fig.S4). This is consistent with a higher selective pressure in larger populations. Therefore, when the cost of immunity is high, the immune response can be downregulated after only two signaling steps (*R* → *A* and *A* → *I*) (Fig.4c2). In contrast, when the cost of immunity is low, downregulation occurs after three steps (*R* → *A, A* → *D*, and *D* → *I*) (Fig.3).

**Fig 3.**
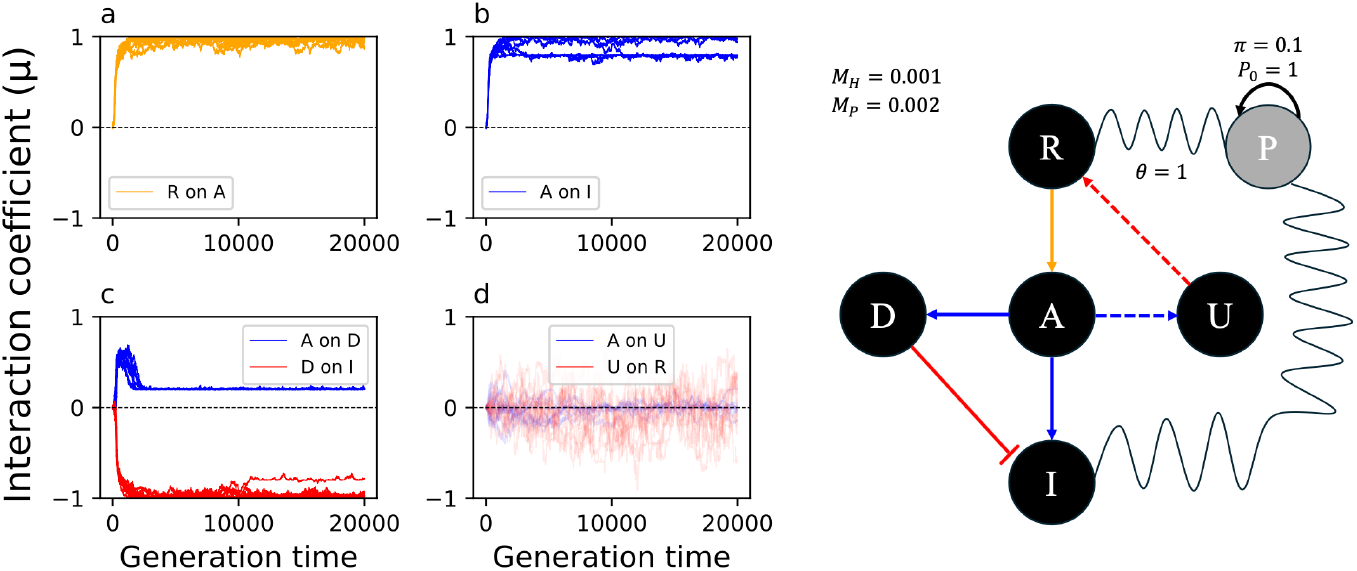
The evolution of immune signaling when the protein degradation rate is zero. The Y-axis in the left panel shows the coefficient of interaction (*μ*_*i,j*_) from −1 (complete inhibition) to +1 (complete activation). The interaction does not evolve when *μ* fluctuates around 0. The X-axis shows the generation time. The right panel shows the evolved signaling pathway after 20,000 generations. The color of the arrows in the right panel corresponds to the color of the plots in the left panel. The oscillating lines represent co-evolution between the host and the pathogen. The dashed arrows show the interactions that did not evolve. Here *θ* = 1; thus, all individuals are infected. The initial condition (*P*_0_) for the pathogen, its replication rate (*π*), and mutation rate for the host (*M*_*H*_) and pathogen (*M*_*P*_) are shown in the right panel.

**Fig 4.**
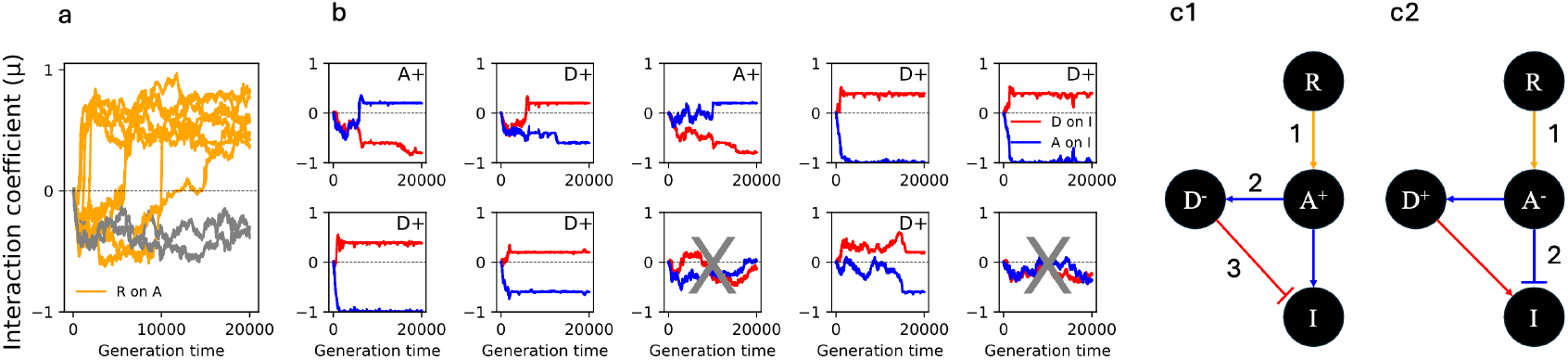
A high cost of immunity results in the evolution of two types of signaling networks with a downstream NFL. Panel a shows whether the first signaling step (activation of *A* by *R*) evolves. The Y-axis shows the coefficient of interaction (*μ*) and the X-axis is the generation time. The plots in panel b (10 plots showing 10 replicate simulations) show the effect of *D* (red) and *A* (blue) on *I*. In two simulations, the activator (*A*) activates the immune response and the downstream regulator (*D*) acts as an NFL. These are annotated as *A*+. In six simulations the downstream regulator activates immunity (*D*+) and the activator protein (*A*) functions as an NFL. In two simulations the receptor does not activate the activator protein (grey trajectories in panel a); thus, signaling is not transmitted to the host proteins. These two simulations are marked with grey crosses in panel b. The two types of evolved signaling networks are shown in panels c1 and c2, where the numbers indicate the number of steps needed to downregulate *I*.

### The evolution of the upstream NFL is influenced by the protein degradation rate, receptor-pathogen coevolution, infection rate, and population size

In previous simulations, we observed that setting *ϕ* > 0 led to some activation of *U* (the upstream regulator) by *A*; however, *U* still did not downregulate the immune response (Fig.S2). When *ϕ* > 0, the equilibrium concentration of host proteins depends on their degradation rate and activation by an upstream protein. To further enhance the transfer of information from upstream to downstream proteins, we set *μ*_*R,P*_ (Fig.2) to 1. This ensures that the receptor doffs not co-evolve with the pathogen and is fully activated by it. This might reflect a real-world situation where the receptor recognizes cytokines or microbial structures that are evolutionarily conserved. Under these conditions, the upstream NFL (*U*) evolved in nine out of 10 simulations (Fig.5a). When we set the degradation rate of the receptor to 0 (*ϕ*_*i*≠*R*_ > 0, *ϕ*_*i*≠*R*_ = 0, and *μ*_*R,P*_ = 1), the upstream NFL (*U*) evolved in all 10 simulations (Fig.5b). In the absence of pathogen-receptor co-evolution, the target of the upstream NFL (*R*) exclusively evolves with *U*, whereas the target of the downstream NFL (*I*) evolves with both *D* and *A*. To reduce this bias while ensuring the effective transfer of information from the pathogen to the receptor, we set the mutation rate of the pathogen to zero (M_*P*_ = 0) while allowing the receptor to evolve and detect the pathogen. As before, the upstream NFL (*U*) evolved to reduce the cost of the immune response (Fig.S5).

We next examined the effect of the rate of infection on the evolution of the upstream NFL. To this end, we reduced the infection rate (*θ*) from 1 to 0 .7 in a population of 2,000 individuals. For this simulation, we chose the parameter values under which both *U* and *D* previously evolved as NFLs (*ϕ*_*i*≠*R*_ > 0, *ϕ*_*i,R*_ = 0, and *μ*_*R,P*_ = 1). We observed that in a minority of the simulations, *U* did not evolve as an NFL (left panel of Fig.5c). We hypothesized that under a smaller infection rate than 1, there is less evolutionary pressure to reduce the cost of infection. This is because uninfected parents would survive and contribute to the offspring of the next generation; thus, resulting in a larger effect of drift. To test this, we increased the population size from 2, 000 to 5,000 and repeated the simulation with *θ* = 0 .7. As expected, in all simulations both *U* and *D* evolved as NFLs to reduce the cost of immunity (right panel of Fig.5c).

Next, we asked how increasing the cost of immunity (setting *α* = 2 and *β*= 2 in *Eq. 4*) would affect the evolution of the upstream NFL under the condition in which it previously evolved (*ϕ*_*i*≠*R*_ > 0, *ϕ*_*i*=*R*_ = 0, and *μ*_*R,P*_ = 1). In some simulations, the upstream NFL did not evolve when the cost of immunity was high (*β*= 2) (left panel of Fig.5d). In these simulations the activation of the immune response by *A* was reduced (right panel of Fig.5d). This leads to a more direct control of *A* on *I* without the involvement of *U* as an NFL. Thus, a higher cost of immunity does not guarantee the evolution of an upstream NFL.

**Fig 5.**
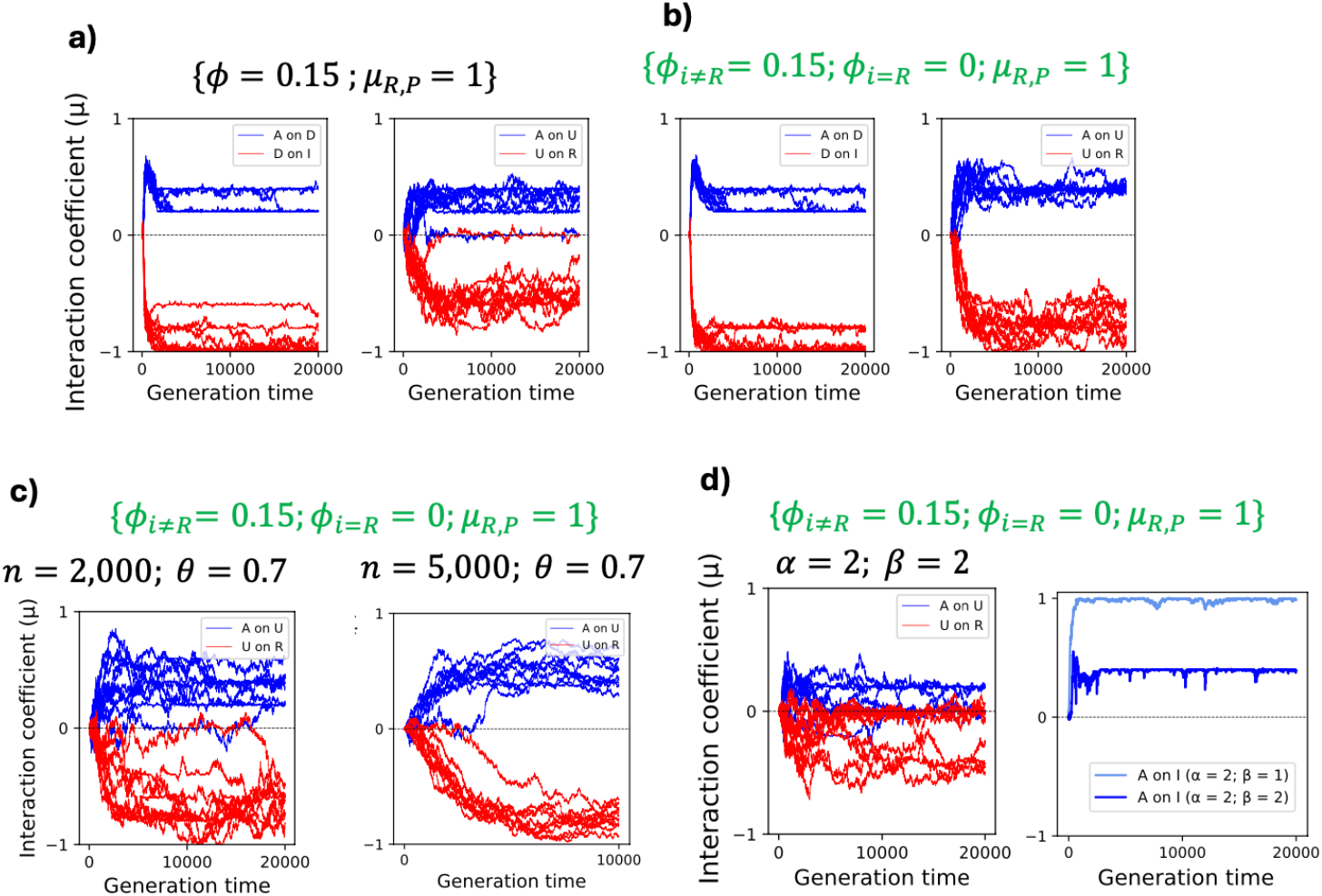
The evolution of the upstream NFL is sensitive to changes in the parameter values. In panel a, non-zero degradation rate for the host protein (*ϕ* = 0.15) and the absence of co-evolution between the receptor and pathogen (*μ*_*R,P*_ = 1) give rise to the evolution of the upstream NFL in most simulations along with the downstream NFL. In panel b, a more robust evolution of the upstream NFL is observed when amongst the host proteins only the receptor degradation rate is set to 0 (*ϕ*_*i*, ≠ *R*_ = 0.15 and *ϕ*_*R*_ = 0). This condition is shown in green and is repeated in panels c and d. Panel c shows the combined effect of infection rate (*θ*) and population size (*n*). When the infection rate is less than 1 (*θ* = 0.7), the upstream NFL does not evolve in some simulations. Increasing the population size (*n* = 5,000) ensures the evolution of the upstream NFL even when *θ* = 0.7. In panels a-c we set *α* = 2 and *β* = 1. Panel d shows the effect of the cost of the immune response under the conditions that favor the evolution of the upstream NFL. When the cost of immunity is high (*β* = 2) the upstream NFL does not evolve in some simulation. Instead, the effect of *A* on *I* is reduced to attenuate the immune response. In all simulations presented here, *D* evolves as an NFL.

### Estimates of the rate of evolution confirm the robust evolution of the downstream NFL while showing a weak selection on the upstream NFL

We next asked if the selective pressures acting on upstream and downstream NFLs in our model are consistent with the general distribution of the observed *ω* values in Fig.1, as well as the prediction of the model that the evolution of downstream NFLs, unlike the upstream one, is robust to the parameter values. We estimated the rate of evolution of host proteins by comparing the rate of change in the functional domains (sender and receiver) to the neutral domain (*Eq. 7*), when the rate of change for the neutral domain is below a threshold (Fig.6a). We estimated the global rate of evolution for all host proteins as a reference (Fig.6b) to evaluate the size effect of *ω* for the upstream (*U*) and downstream (*D*) regulators (Fig6.c). To this end, we chose three of our original scenarios: 1) parameters in Fig.3 in which *D* evolves as an NFL (purple trajectories in Fig.6c), 2) parameters in Fig.4 which give rise to polymorphism, and either *A* or *D* evolves as the downstream NFL (yellow trajectories in Fig.6c), and 3) parameters in Fig.5b under which both regulators (*D* and *U*) evolve as NFLs (cyan trajectories in 6c). These scenarios encompass all types of NFLs that evolved in our simulation (only downstream: *D* or *A*, and both upstream and downstream: *U* and *D*). We found that when *D* evolves as an NFL both receiver and sender domains of *D* are under negative selection (purple and cyan trajectories in the left panels of Fig.6c). On the other hand, there is a weak evolutionary pressure on *D* when there is polymorphism in the evolution of the downstream NFL (yellow trajectories in left panels of Fig.6c). This is consistent with replaceable function of *D* by *A* as an NFL. Consistent with previous predictions of the model, across the three simulations, *ω* for *U* fluctuates around 0, and the average *ω* is very small (right panels of Fig.6c) given its global distribution (Fig.6b). This indicates weak selective pressure on *U*.

**Fig 6.**
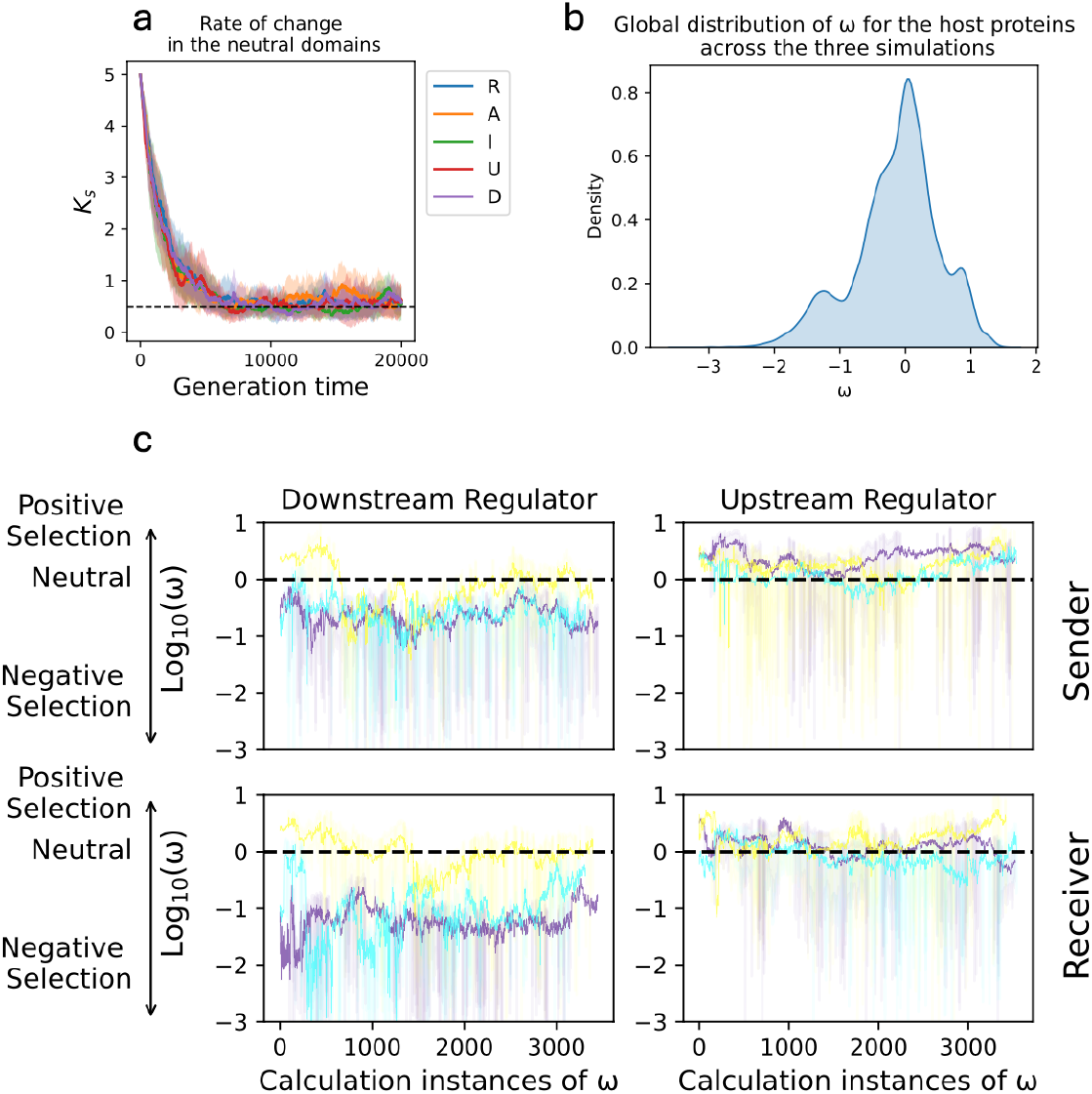
The downstream regulator evolves under conservative evolution, whereas the upstream regulator is under weak selective pressure. Panel a shows a typical rate of change for the neutral domain (*k*_*s*_) of all host proteins across simulations. Panel b shows the distribution of pooled *ω* values across three simulations for all proteins of the host. Panel c shows the evolution of downstream and upstream regulators. The rate of evolution (*ω*) is calculated every five generations and only when *k*_*s*_ < 0 .5. The X-axis shows the instances of calculation *ω* through the evolution, and the Y-axis shows the rate of evolution for the two domains (rows) of *D* (left column) and *U* (right column). The trajectories show the average rate of evolution across 10 simulations. The shaded regions specify the standard deviation. Positive values on the Y-axis show positive selection, 0 indicates neutral evolution and negative values specify negative selection. The evolution of *D* and *U* are shown under three parameter regimes. Under one regime *D* evolves as an NFL (purple). Under the second regime, there is polymorphism in the choice of the downstream NFL (either *A* or *D*), which is shown by the yellow trajectories. The cyan trajectories specify the parameter regime under which both *U* and *D* evolve as NFLs.

## Discussion

We modeled the evolution of immune signaling and uncovered distinct patterns of evolution for upstream and downstream NFLs consistent with the observed rate of evolution (Fig.1). We found that the evolution of a downstream NFL is robust to changes in the parameters; however, the exact architecture of a signaling pathway with a downstream NFL is linked to the cost of the immune response. Specifically, we found that a higher cost of immunity results in a shorter path to inhibition of the immune response and a longer path to its activation (Fig.4). We also found that the evolution of an upstream NFL is contingent upon the absence of host-pathogen co-evolution, degradation of signaling proteins, and a high infection rate (Fig.5). Given the ubiquitous phenomenon of multi-level regulation within signaling pathways, this study is a significant step in understanding regulation mechanisms across diverse signaling pathways.

### The evolution of the downstream NFL is robust to changes in model parameters

Our model predicts that the evolution of a downstream NFL is robust to factors such as the degradation rate of signaling proteins, pathogen replication rate, rate of infection, population size, and the cost of immunity (Fig.3 and Fig.S2). Downstream NFLs across various signaling pathways show a slower and more consistent rate of evolution compared to upstream NFLs (Fig.1). Downstream NFLs, unlike upstream ones, are not dependent on signaling proteins, and directly regulate the expression of target genes, making them more effective in reducing the cost of signaling. Based on our model predictions, we propose that a slow rate of evolution for downstream NFLs could result from the effectiveness of these NFLs in adjusting the output of signaling pathways without the involvement of intermediate components.

Downstream regulators of immune signaling have been identified across many signaling pathways. The Toll pathway, which is one of the primary immune signaling pathways in invertebrates, is regulated by two redundant downstream regulators Cactus and WntD, which act as downstream NFLs by inhibiting the transcription factors Dif/Dorsal responsible for inducing immune genes (Aggarwal & Silverman, 2008). On the other hand, even though there are constitutive negative regulators upstream of Toll signaling in *Drosophila*, no upstream NFLs have been identified that uniquely regulate this pathway. The robust evolution of downstream NFL in our model as opposed to the parameter-sensitive evolution of upstream NFL shed light on such differences in the use of NFLs within biological networks. In addition, the robust evolution of downstream NFLs in our model aligns with the consistency in the mechanisms of action of downstream NFLs (e.g., Repressosome, IκBα, and JAM) across various immune signaling pathways (Kearns et al., 2006; Kim et al., 2007; Sasaki-Sekimoto et al., 2013). We propose that a direct interaction between downstream NFLs and target genes leads to the evolution of similar downstream NFLs across various signaling networks.

### The cost of immunity determines the architecture of the signaling network with a downstream NFL

We found that when the cost of the immune response is high, the number of steps leading to the inhibition of the target gene is shorter than those for its activation (Fig.4). Previous studies focused on the evolution of complexity —such as the number of proteins and their interactions within signaling pathways— without distinguishing between steps required for inhibition or activation (Lynch, 2007; Soyer & Bonhoeffer, 2006). Addressing this distinction is crucial because signaling pathways are regulated by NFLs, which are either transcriptionally induced or activated through post-translational modification of existing proteins. For example, ERK1/2 MAPK pathway, which regulates fundamental cellular processes such as proliferation, differentiation, and survival is regulated by multiple NFLs that are either induced transcriptionally, such as DUSP, and by direct phosphorylation of signaling proteins (Lake et al., 2016). In the ERK1/2 MAPK pathway, the downstream protein ERK1/2 can inhibit its own activator MEK 1/2 via phosphorylation (Lake et al., 2016). This reduces the number of signaling steps necessary to inhibit gene expression compared to the induction of the gene encoding for DUSP. Our model predicts that a high cost of signaling, which is typical of pathways such as ERK1/2 MAPK that control fundamental cellular processes, can result in polymorphism in the evolution of downstream NFLs. These predictions are derived, of course, from a minimal model that does not capture the nuances of different regulation mechanisms such as transcriptional induction of NFLs and regulation via phosphorylation. Future studies that specifically model multiple mechanisms of NFLs can shed more light on the evolution of such regulatory motifs within biological networks.

### The evolution of the upstream NFL is sensitive to the degradation rate of signaling proteins, host-pathogen co-evolution, infection rate, and population size

The large variation in empirically observed evolutionary rates amongst upstream NFLs (Fig.1) suggests that their position within signaling pathways does not strongly predict evolutionary rate. This implies that, unlike downstream NFLs, upstream NFLs may not be under strong selective pressure. Consistent with this hypothesis, our model predicts that the evolution of the upstream NFL occurs only under specific conditions because they play a less critical role in regulating target genes compared to their downstream counterparts in most situations.

We found that a non-zero degradation rate of intracellular signaling components (excluding *R*) is necessary but not sufficient for the evolution of the upstream NFL (Fig.5a). This is because proteins that are inactivated by degradation over time rely on upstream signals to sustain a certain level of activity within the signaling pathway. Thus, a non-zero degradation rate creates a dependency between upstream and downstream components. Without this dependency, an NFL cannot reduce the cost associated with the production of downstream components (e.g., *I*) by inhibiting upstream proteins. Within signaling pathways, protein degradation serves multiple purposes including the removal of damaged and misfolded proteins (Goldberg, 2003), cell cycle progression (Dang et al., 2021), and facilitation of signaling (e.g., degradation of IκBα activates NF-κB) (Mathes et al., 2008). We propose a new role for protein degradation within signaling pathways: it establishes a stronger hierarchy of protein-protein interactions, allowing the upstream NFLs to reduce the cost of response. In addition, we found that a zero degradation rate for the receptor facilitates the evolution of the upstream NFL (Fig.5b). This is consistent with the role of NFLs such as Pirk in Imd signaling, which removes cell surface receptors rather than relying on their natural degradation (Lhocine et al., 2008).

In addition to a non-zero degradation rate for signaling proteins, we found that the absence of co-evolution between the host receptors and the pathogen is necessary for the evolution of an upstream NFL (Fig.5a). This is because, in the absence of co-evolution, the receptor is fully activated by the pathogen and transmits the information to downstream proteins, establishing a strong link between upstream and downstream components. As a result, the upstream NFL can reduce the cost of the downstream immune response by targeting upstream components. In addition, the upstream NFL would be less effective if it interacts with the constantly changing domain of the receptor that co-evolves with the pathogen. In nature, upstream NFLs such as A20 and Pirk in NF-κB signaling pathway interact with the cytoplasmic domain of the receptor, which does not co-evolve with the pathogen (Kleino et al., 2008; Pujari et al., 2013). In addition, these immune pathways are often activated by conserved pathogen-associated molecular patterns or by cytokines (Li & Wu, 2021). Such interactions minimize rapid changes in response to co-evolution. Examples from signaling pathways where the receptor does directly interact with the pathogen can help reinforce this intuition. For example, type I interferon signaling is initiated by an interaction between the receptor MDA5 and RNA viruses (Dias Junior et al., 2019). This interaction activates an upstream NFL involving the cleaved form of 14-3-3η (sub-14-3-3η) that inhabits MDA5 (Chan et al., 2024). MDA5 indeed co-evolves with viruses (Geng et al., 2024), and human coronavirus can promote the activation of the NFL to reduce MDA5 activity (Chan et al., 2024). Given these observations, it is surprising that MDA5 is regulated by an NFL. However, MDA5 interacts with RNA virus via the C-terminal domains (helicase domain and CTD) and is downregulated by sub-14-3-3η through the N-terminal domain (CARD) (Dias Junior et al., 2019; Lin et al., 2019). This separation of functions across multiple domains prevents host-pathogen co-evolution from affecting the interaction between the receptor and NFL. We propose that such functional division across multiple domains is crucial for the maintenance of upstream NFLs such as sub-14-3-3η.

Consistent with the large variance in 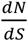 values for genes encoding upstream NFLs (Fig.1), our model predicts a weaker selection pressure on the upstream (*U*) regulator compared to the downstream (*D*) regulator (Fig.6). This is manifested in two of our findings. Firstly, our model predicts that if the infection rate is low and not all individuals in the population are infected, the upstream NFL is less likely to evolve (left panel of Fig.5c). However, in larger populations, where drift has less influence, the upstream NFL evolves even when the infection rate is below 1 (right panel of Fig.5c). Our study is the first to consider the effect of population size on the evolution of NFLs. Previous studies have shown that bacterial proliferation rate and the frequency of encounters with pathogens can significantly affect the optimization of upstream NFLs; however, these models did not consider the evolution of NFLs across generations (Asgari et al., 2024; Asgari & Tate, 2024). Here, we propose that if drift is strong, such factors may have less influence on the evolution of upstream NFLs. We also found that a high cost of immune response does not guarantee the evolution of an upstream NFL (Fig.5d). This happens because an urgent need for lowering the immune response is more likely to be regulated by downstream components that directly interact with immune genes. In our model, this is reflected in the weaker activation of the immune response by the central activator. Knock-down studies of NFLs have shown that NFLs reduce the cost of signaling (Beg et al., 1995; Kim et al., 2007; Prakash et al., 2024). Thus, our finding challenges the well-established view that the evolution of NFLs is directly linked to the cost of signaling.

## Conclusions

Our model provides novel insights into the evolution of NFLs functioning at different signaling stages. Consistent with the more conservative rate of evolution for downstream NFLs (Fig.1) we found these NFLs evolve robustly under most parameter and co-evolutionary scenarios, signifying their importance in the regulation of signaling pathways. In addition, our model proposes that the following is necessary for the evolution of upstream NFLs: 1) degradation of proteins transmitting the signal, 2) lack of co-evolution between receptor and pathogen, and 3) large number of infected individuals per generation (high rate of infection or lower rate of infection in larger populations). We believe that such insights significantly advance our understanding of the evolution of signaling pathways amongst diverse species.

## Materials and Methods

### The model

Our model captures fundamental elements common to immune signaling networks across a variety of taxa (Fig. 2). In this model, each host encounters a pathogen (*P*) with a probability *θ*, and the pathogen proliferates at a rate *π*. Signaling is triggered by an interaction between the pathogen and the receptor (*R*). The receptor subsequently engages with the central activator (*A*) protein. Next, *A* interacts with the immune (*I*) protein and two regulator proteins. One regulator functions upstream (*U*) of the pathway and interacts with *R*; thus regulating the input into the pathway. The other functions downstream (*D*) to regulate the output of signaling (*I*). This model design allows us to explore the evolution of an immune signaling network (*R* → *A* → *I*) that responds to the pathogen and is modulated by an upstream (*U*) and a downstream (*D*) regulator.

Kamiya et al. (2016) used a minimal evolutionary model to study the effect of host-pathogen co-evolution on investment in induced and constitutive defenses. Following their approach, we simulated the signaling proteins and pathogen with three domains (sender, receiver, and neutral), each containing *L* = 10 bits. The receiver domain of each protein (*i*) interacts with the sender domain of another protein (*j*) according to the network in Fig.2. The neutral domain does not participate in protein-protein interactions, serving instead as a benchmark for the rate of evolution. The coefficient of interaction between two proteins (*μ*_*i,j*_) is determined by the hamming distance, which is the total number of bits that are different at corresponding sites between the sender domain of the acting protein and the receiver domain of the target protein (*Eq. 1*). When the number of matching positions between the two domains exceeds 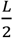, the interaction is activatory (1 ≥ *μ*_*i,j*_ > 0); if fewer than 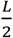, the interaction is inhibitory (−1 ≤ *μ*_*i,j*_ < 0). If there are exactly 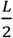 matches, no interaction takes place (*μ*_*i,j*_ = 0). The only interaction forced to evolve as inhibitory is between immunity (*I*) and the pathogen (*P*) (*Eq*. 2). We modeled changes in the concentration of the host proteins over the host lifetime using a system of ODEs, following Kamiya et al. (2016) (*Eq*. 3). The host proteins degrade at a rate *ϕ*. Studies on immune signaling have shown that protein degradation increases following signaling (Marchingo & Cantrell, 2022) and central transcription factors, such as NF-κB in its active state, have a relatively short half-life (Bergqvist et al., 2009; Hohmann et al., 1991), as do their inhibitors (Hay et al., 1999). This suggests a non-zero value for *ϕ* during the signaling period. In our simulations, we check for both *ϕ* = 0 and *ϕ* > 0. We assume that the pathogen does not spontaneously degrade. This promotes the evolution of a functional immune signaling network that fights oE pathogens. If the interaction is activatory, the concentration of the acting protein *y*_*j*_ is multiplied by one minus the concentration of the target protein *y*_*i*_. This ensures that the concentration of each protein including the pathogen is between 0 and 1. For the infected host, the initial condition of the pathogen (*P*_0_) is 1, unless otherwise specified. The initial condition for host proteins is 0. The lifespan of the host is *T* = 1,000.

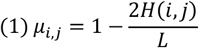

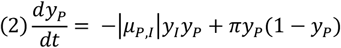

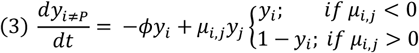

### Evolutionary simulations

We ran evolutionary simulations to identify conditions under which a functional signaling network evolves (*R* activates *A* and *A* activates *I*), which is negatively regulated by upstream and downstream NFLs (*μ*_*R,U*_ < 0 and *μ*_*I,D*_ < 0). Each simulation starts with *n* randomly created haploid hosts and pathogens. Unless otherwise specified, *n* = 2,000 for both hosts and pathogens and the number of generations is 20,000. In each generation, a host encounters a pathogen randomly. Hosts reproduce sexually while pathogens reproduce asexually. During sexual reproduction, for each host protein, the domain and the site within that domain where recombination occurs are selected at random. One recombination is allowed per protein during reproduction. The sequence to the left of the recombination site comes from one parent and the sequence to the right of that site comes from another parent. A domain that does not recombine is randomly inherited from either of the two parents. The hosts mutate at a rate M_*H*_ = 0.001 in a randomly chosen protein, domain, and site. Because pathogens often mutate faster than the host (Hafner et al., 1994), in our simulations, pathogens mutate at twice the host mutation rate (M_*P*_ = 2 M_*H*_), unless otherwise specified. The parents and pathogens are chosen for reproduction based on their relative fitness in the population. The fitness of an infected host is modeled as an exponential function which is reduced from 1 to 0 as the average concertation of the pathogen and immunity (due to energetic costs and immunopathology) increases over the host lifetime (*Eq. 4*). The coefficients *α* and *β*in the host fitness function represent the relative costs of infection and immune response, respectively. The fitness of a pathogen infecting a host depends on its average concentration through the lifetime of the host (*Eq. 5*). We do not assume constitutive investment in immunity; thus, the uninfected host has a fitness of 1. When the infection rate is less than 1, some pathogens, by random chance, fail to infect hosts. These pathogens then have a fitness of 0. For every parameter combination, we ran a total of 10 independent replicate simulations.

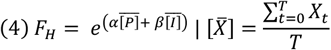

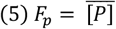

To distinguish evolutionary forces that act upon the upstream (*U*) and downstream (*D*) regulators, we estimated the rate of evolution in sender and receiver domains following Kamiya et al. (2016). To this end, every five generations, we calculated the consensus sequence, which is the average proportion of 1s across all sites within the population. If *t* = 0, the consensus sequence is referred to as the focal consensus (*FO*); otherwise, it is referred to as the temporal consensus (*TE*). The rate of change for a neutral domain with *L* = 10 bits follows *Eq. 6*. The first term in the sum specifies the proportion of changes from 1 to 0 and the second term changes from 0 to 1:

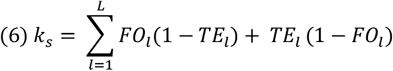

If *k*_*s*_ exceeds 0 .5 (saturation of the neutral domain), we set *FO* = *TE*, otherwise we calculate the rate of evolution for sender and receiver domains (with a changing rate *k*_*a*_) using *Eq. 7*:

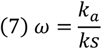

## Supporting information

Supplemental file for the manuscript titled How the Structure of Signaling Regulation Evolves: Insights from an Evolutionary Model

## Acknowledgments

This work was supported by the National Institute of General Medical Sciences at the National Institutes of Health (grant number R35GM138007 to A.T.T.).

## Author Contributions

D.A. and A.T.T. conceived and designed the study. D.A. performed the simulations and wrote the first draft. D.A. and A.T.T. contributed to the final draft.

## Data Availability Statement

We performed all simulations in Python. The code repository is accessible at: https://github.com/danialasg74/NETWORKEVOLUTION/blob/main/code.py

