## Supplemental file for the manuscript titled How the Structure of Signaling Regulation Evolves: Insights from an Evolutionary Model for "How the Structure of Signaling Regulation Evolves: Insights from an Evolutionary Model"

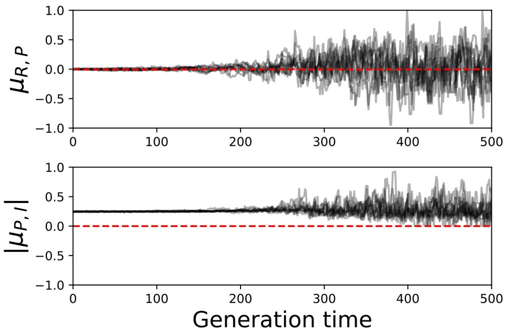


**Fig.S1. Co-evolution between the receptor and pathogen (upper panel) and between the immune protein and pathogen (lower panel).** The fluctuations in the coefficient of interactions represent the continuous co-evolution between the host and parasite. The absolute value of the coefficient of interaction between the immunity and pathogen is shown here. In the system of ODE this value is multiplied by $-1$; thus, the effect of immune response on the pathogen solely evolves as inhibitory.

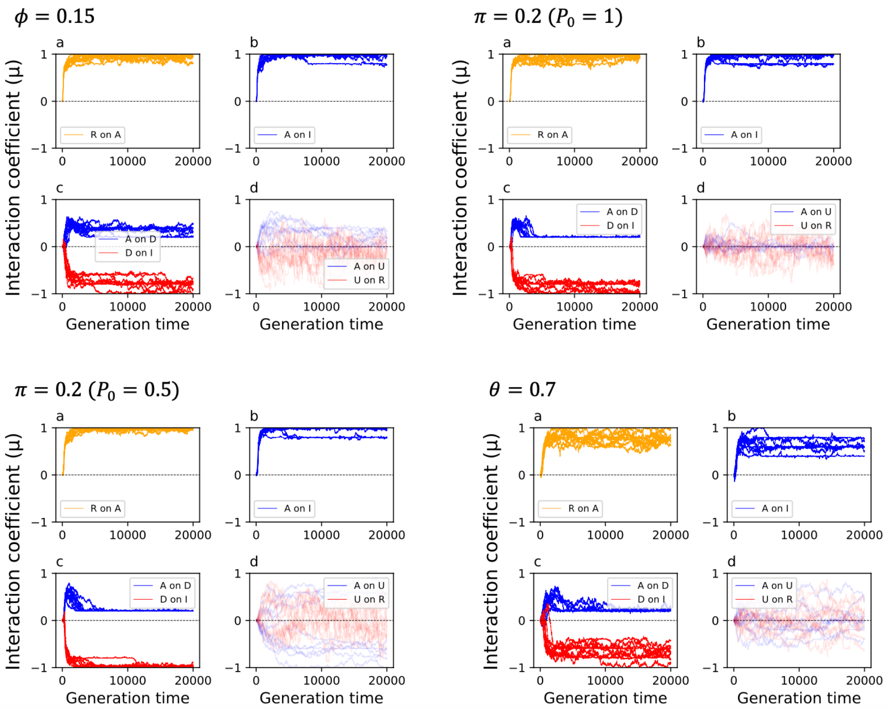


**Fig.S2. The evolution of the downstream NFL is robust to the choice of parameters.** Here we show the effect of the host protein degradation rate (top left), pathogen proliferation rate (top right), initial condition of the pathogen and its proliferation (bottom left), and the rate of infection (bottom right). In all cases, a functional signaling pathway evolves (a and b panels), which is regulated by a downstream NFL (c panels). However, the upstream NFL does not evolve (d panels).

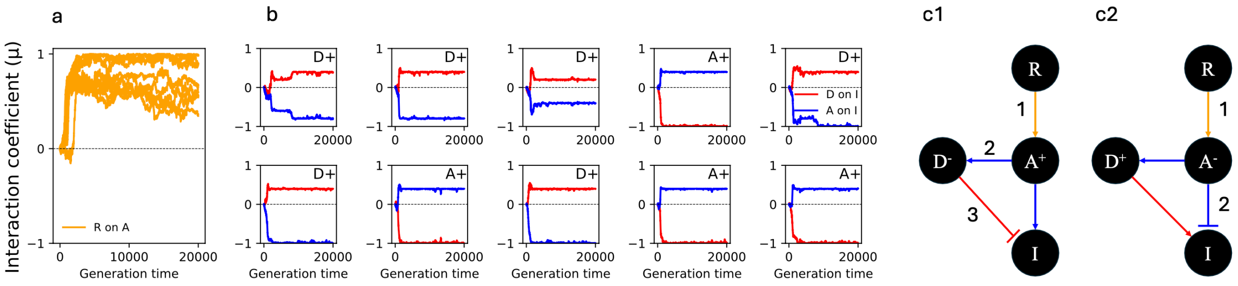


**Fig.S3. The evolution of polymorphism in the architecture of immune signaling due to the high cost of immune response is robust to host protein degradation rates.** Here we set $\phi = 0.15$ and other parameter values are identical to those used for Fig.3 in the main text. Panel a shows whether the first signaling step (activation of $A$ by $R$) evolves. The Y-axis shows the coefficient of interaction ($\mu$) and the X-axis is the generation time. The plots in panel b show the effect of $D$ (red) and $A$ (blue) on $I$. In four simulations, the activator ($A$) activates the immune response and the downstream regulator ($D$) acts as an NFL. These are annotated as $A$+. In the remaining six simulations the downstream regulator activates immunity ($D$+) and the activator protein ($A$) function as an NFL. The two types of evolved signaling networks are shown in panels c1 and c2. The numbers in panels c1 and c2 show the number of steps needed to downregulate $I$.

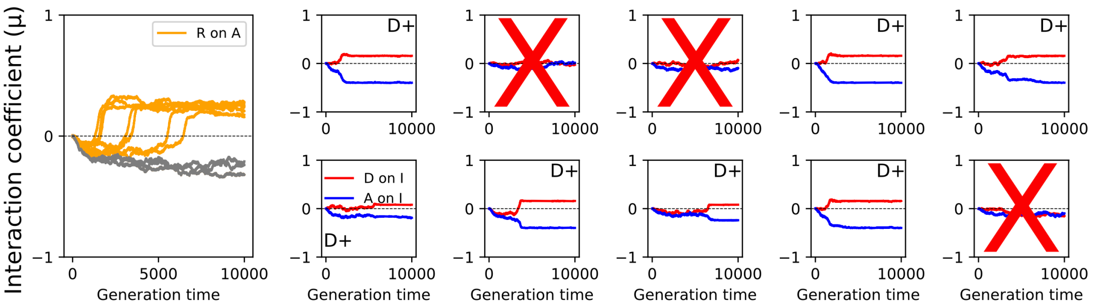


**Fig.S4. A large population size ($n = 5,000$) results in the evolution of $A$ (blue lines) as an NFL in all simulations if the cost of the immune response is high ($\beta = 2$).** In three simulations, the signaling network does not evolve. These are shown as grey trajectories in the left panel and marked by red crosses in the right panels.

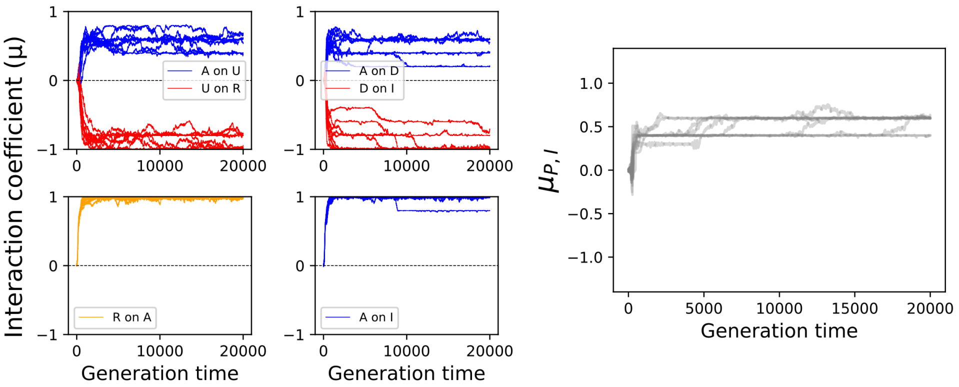


**Fig.S5. Setting the mutation rate of the pathogen to 0 results in the evolution of the upstream NFL.** Here the receptor evolves with the pathogen to activate the immune response (the right panel).
